# JNK activation dynamics drive distinct gene expression patterns over time mediated by mRNA stability

**DOI:** 10.1101/2025.05.02.651814

**Authors:** Abbas Jedariforoughi, Rachel Burke, Andrew Chesak, Jose L. Gonzalez Hernandez, Ryan L. Hanson

**Author notes:** denotes equal contribution (Abbas Jedariforoughi, Rachel Burke).

## Abstract

c-Jun N-terminal kinase (JNK) plays a major role in the regulation of cell death. Numerous studies have highlighted how the dynamics of this kinase dictate whether cells survive in response to cellular stress or induce cell death mechanisms. However, it is less clear how these dynamics potentially contribute to downstream gene expression patterns through regulated transcription factors like c-Jun. To investigate this question, we used a treatment strategy with the JNK agonist anisomycin to drive specific dynamics; sustained, transient, or pulsed activation, and assessed the impact on downstream gene expression patterns. We observed that multiple gene expression patterns emerged depending on the dynamics of JNK activation. Ordinary Differential Equation (ODE) models suggest that a subset of these clusters are mediated by mRNA stability and supported by measured mRNA decay rates. Specific gene clusters also show enrichment in specific cellular pathways, including cell death and inflammatory signaling, suggesting these dynamics contribute to differential regulation of these pathways. These findings highlight another contribution of JNK dynamics to the regulation of cellular responses to stress stimuli.

## Introduction

Cells maintain homeostasis by rapidly sensing and responding to internal and external changes in cell state. Cells have several mechanisms to distinguish between different stimuli, for example, different receptors that distinguish between ligands^1-3^, differential activation of specific signaling molecules^4^, as well as differences in the temporal dynamics of the specific downstream pathways^5,6^. This process of “dynamic encoding” can enable cells to distinguish between stimuli that converge upon the same signaling networks. For example, temporal dynamics of the stress-responsive transcription factor p53 can allow cells to distinguish between ultraviolet light (UV)-induced damage, DNA double-strand breaks (DSBs), as well as specific chemotherapies^7-11^. These dynamics can give rise to specific gene and protein expression patterns^7,12-14^, as well as elicit specific cell fate outcomes^8,9^. Similarly, NF-kB exhibits stimulus specific dynamics following exposures to pathogen associated molecular patterns (PAMPs), damage associated molecular patterns (DAMPs), and cytokines to drive unique gene expression patterns^3,15^. The precise gene expression dynamics elicited in the case of p53 and NF-kB rely both on the dynamics of the transcription factor itself, as well as the mRNA stability of the specific target gene^6,7,12^.

Dynamic encoding is not strictly limited to transcription factors, as the mitogen-activated protein kinase (MAPK) c-Jun N-terminal kinase (JNK) has also been proposed to exhibit dynamic encoding as a means to regulate cell fate outcomes in response to cellular stress. Specifically, the distinction between pro-survival and pro-death signaling. Initial characterization of JNKs function in the mid-1990s highlighted the potential role for the duration of JNK activation in mediating cell death or apoptosis in response to UV damage^16^. Similar approaches demonstrated a biphasic JNK responses in mediating pro-survival or pro-death responses following induction of endoplasmic reticulum stress induced by tunicamycin or thapsigargin^17^. More recently, use of fluorescent biosensors of JNK activation^18^ have facilitated the tracking of single cell kinase activity and cell fates. These studies have highlighted roles for JNK dynamics in regulation of cell death in response to UV^19^, oxidative stress^20^, and activation of the inflammasome in pyroptosis^21^.

While these studies have highlighted clear roles of JNK dynamics in mediating cellular responses, the downstream regulatory impacts of these dynamics remain less clear. Unlike the transcription factors p53 and NF-kB, JNK exerts its effects by phosphorylation of serine and threonine residues on its nearly 100 substrates^22,23^. These include non-nuclear substrates Bcl2, Bax, Bim, and Bad which may regulate cell fate independent of new gene expression. However, in addition to these non-nuclear substrates JNK is also known to phosphorylate several transcription factors to regulate downstream gene expression patterns. The most-well characterized of these substrates is c-Jun, which is phosphorylated on the N-terminal residues serine-63 and serine-73 to potentiate its transcriptional activity as part of the dimeric transcription factor AP-1^22,23^. Currently, it’s unclear how these dynamic patterns of JNK activation potentially contribute to downstream gene expression patterns.

In order to examine this question, we leveraged live-cell imaging approaches of fluorescent biosensors to track JNK activation within single cells. As JNK dynamics can be significantly heterogeneous between single cells and across treatments, we established a dosing regimen of anisomycin to specifically enrich in specific dynamics, notably the duration and number of pulses, and assess gene expression impacts by RNA-seq. We found that cells exhibit an array of gene expression patterns in response to dynamic JNK activation. These dynamics can partially be explained by mRNA stability similar to p53 and NF-kB, but other complex mechanisms are likely present such as transcriptional cross-talk. Interestingly, certain gene clusters are enriched in specific pathways suggesting that dynamics may allow cells to diversify function over time in response to JNK activation. These results provide us new insight into how signal transduction through JNK can contribute to diverse outcomes.

## Results

### JNK dynamics are diverse and heterogeneous across treatments

Previous studies using live-cell imaging and western blot approaches have proposed distinct dynamics of JNK activation in response to diverse stimuli. To examine the emergence and heterogeneity of JNK dynamics, we performed live-cell imaging of RPE1-hTERT cells expressing the JNKKTR fluorescent biosensor to track single cell JNK dynamics. We examined JNK activity in response to sorbitol (200 mM), tunicamycin (1 µM), thapsigargin (1 µM), H_2_O_2_ (50 µM), TGF-β1 (10 ng/ml), and TNFα (20 ng/ml) treatment and analyzed 90+ individual cells per condition over 12.5 hours. Sorbitol, thapsigargin, H_2_O_2_, and TNFα treatment all exhibited a visible response within the population with a rapid pulse of activation (Figure 1A). At the single cell level, individual cells showed marked heterogeneity with some cells responding and others not responding at all (Figure 1B). To identify the dynamics that emerge in response to treatments, we identified the number of pulses of JNK activation by identifying local maxima (Figure 1C) as well as the total duration of activation by quantifying the area under the curve above an assigned threshold of 50% above baseline (Figure 1D). This baseline was established for each condition based on the average of 4 frames before addition of treatments. Overall, only TNF treatment resulted in significant increases in the number of JNK pulses and activity over the 12.5 hour imaging experiment. However, this is likely due to the fact that many treatments induced a singular pulse of activation followed by inactivation. Consistent with this, we observed that sorbitol, thapsigargin, and TNFa all showed significantly longer durations of JNK activity within the first three hours (Figure 1G). Overall, within these experiments we identify a number of dynamics within single cells including singular pulses, multiple pulses, as well as prolonged or sustained activation (Supplemental Figure 1A-D) across all conditions. Therefore, we wanted to interrogate how these particular dynamics may contribute to downstream gene expression patterns.

**Figure 1.**
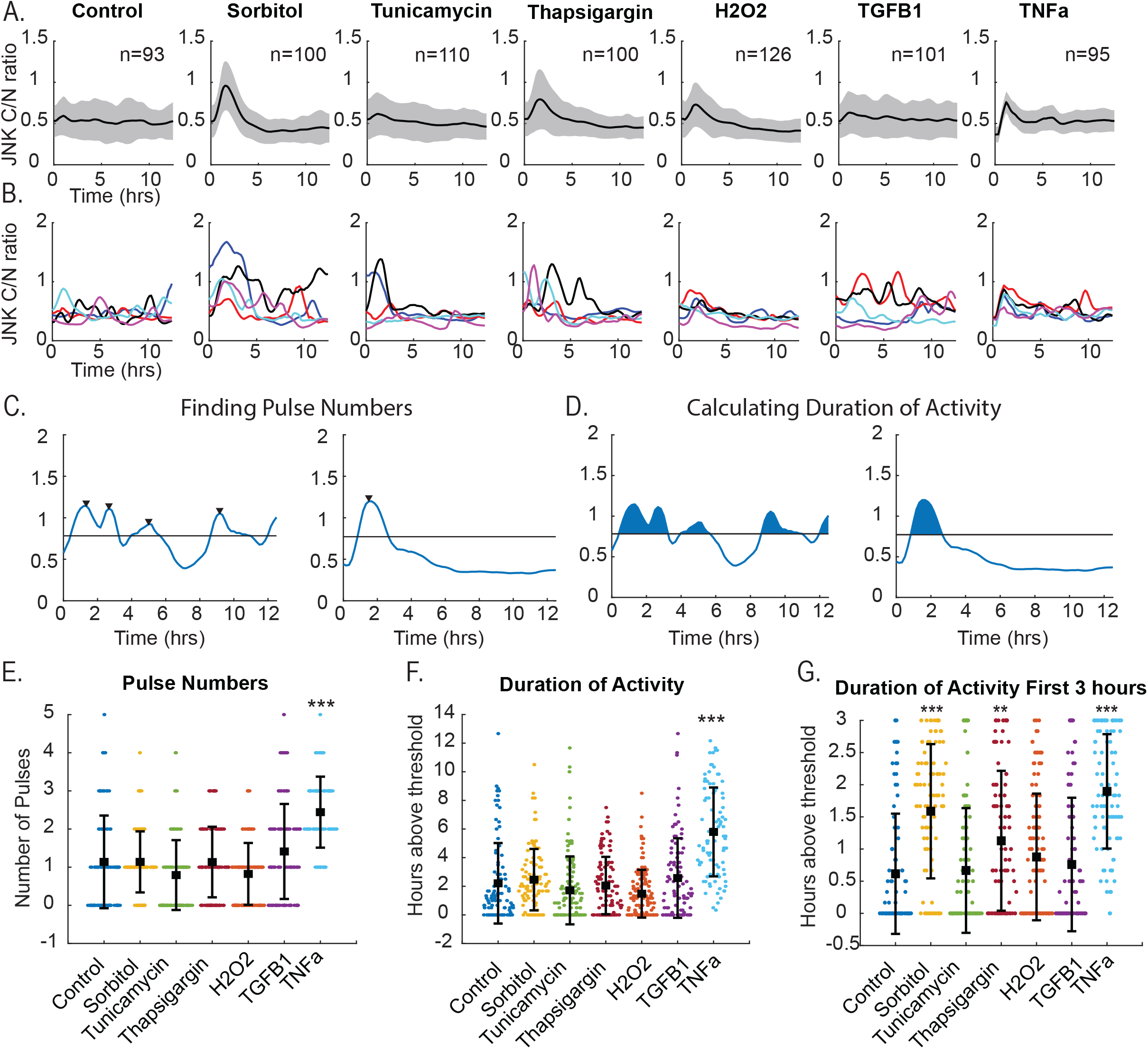
JNK dynamics vary in duration and pulse number across stimuli. **A)** Average cytoplasmic to nuclear ratio of JNKKTR traces of RPE cells treated with DMSO control media, Sorbitol (100mM), Tunicamycin (1 uM), Thapsigargin (1 uM), H_2_O_2_ (50 uM), TGF-β1 (10ng/ml) or TNF (20ng/ml) for 12.5 hours. Thick line represents the average response with the shaded area equal to the standard deviation. n represents the total number of analyzed cells. **B)** Representative cell traces from five individual cells from the same treatment conditions with each line representing the JNK C/N ratio. **C)** Example traces identifying pulse numbers or **D)** total duration of JNK activity within individual cells. **E)** Plot of calculated pulse numbers for each condition. Average and standard deviation for each condition shown. *** p<0.001 as compared to control by ANOVA. **F)** Total duration of activity for each individual cell. Bars show the mean±SD. ***p<0.001 as compared to control by ANOVA. **G)** Duration of JNK activity over the first 3 hours of treatment. Bars represent mean±SD. **p<0.01, ***p<0.001 as compared to control by ANOVA.

### Anisomycin dosing can be used to induce specific dynamic patterns

To study the impact of JNK activation dynamics on gene expression, we decided to enrich in each of the specific dynamics we wanted to examine (transient/singular pulse, multiple pulses, and sustained) as desynchronization within the cell population would mask changes in gene expression over time. To accomplish this, we chose to dose cells with the JNK agonist anisomycin. As anisomycin can inhibit protein translation, we used a subinhibitory concentration of 50 ng/ml to minimize disruptions to protein synthesis^24^. By varying the timing of anisomycin application and removal, we were able to generate three distinct activation profiles: sustained activation, transient activation, and repeated pulses of activation as assessed by live-cell imaging and quantification of JNKKTR cytoplasmic-to-nuclear ratio compared to an untreated control group. Cells were imaged 30 minutes prior to the addition of anisomycin to establish a baseline of activity. In the sustained activation model, continuous exposure to anisomycin led to prolonged JNK activation (Figure 2A). Conversely, transient activation was achieved by adding anisomycin followed by its rapid washout, resulting in a single, brief pulse of JNK activation (Figure 2B). Lastly, repeated pulses of JNK activation were generated by alternating the addition and washout of anisomycin (Figure 2C). At the single cell level, this mechanism of repeated dosing resulted in strong synchronization of the two pulses (Figure 2D). To confirm the anisomycin dosing was driving different dynamics as expected, we calculated overall pulse numbers and total duration of JNK activation over the full 8.5-hour imaging experiment using the same approach as in Figure 1. As expected, we observed that the pulsed cells primarily exhibited two clear pulses and on average more than the single pulse condition as expected (Figure 2E). Furthermore, the sustained condition as expected showed longer durations of JNK activation (Figure 2F). Over the first three hours the transient and pulsed conditions were largely identical and showed no significant difference as expected (Figure 2G).

**Figure 2.**
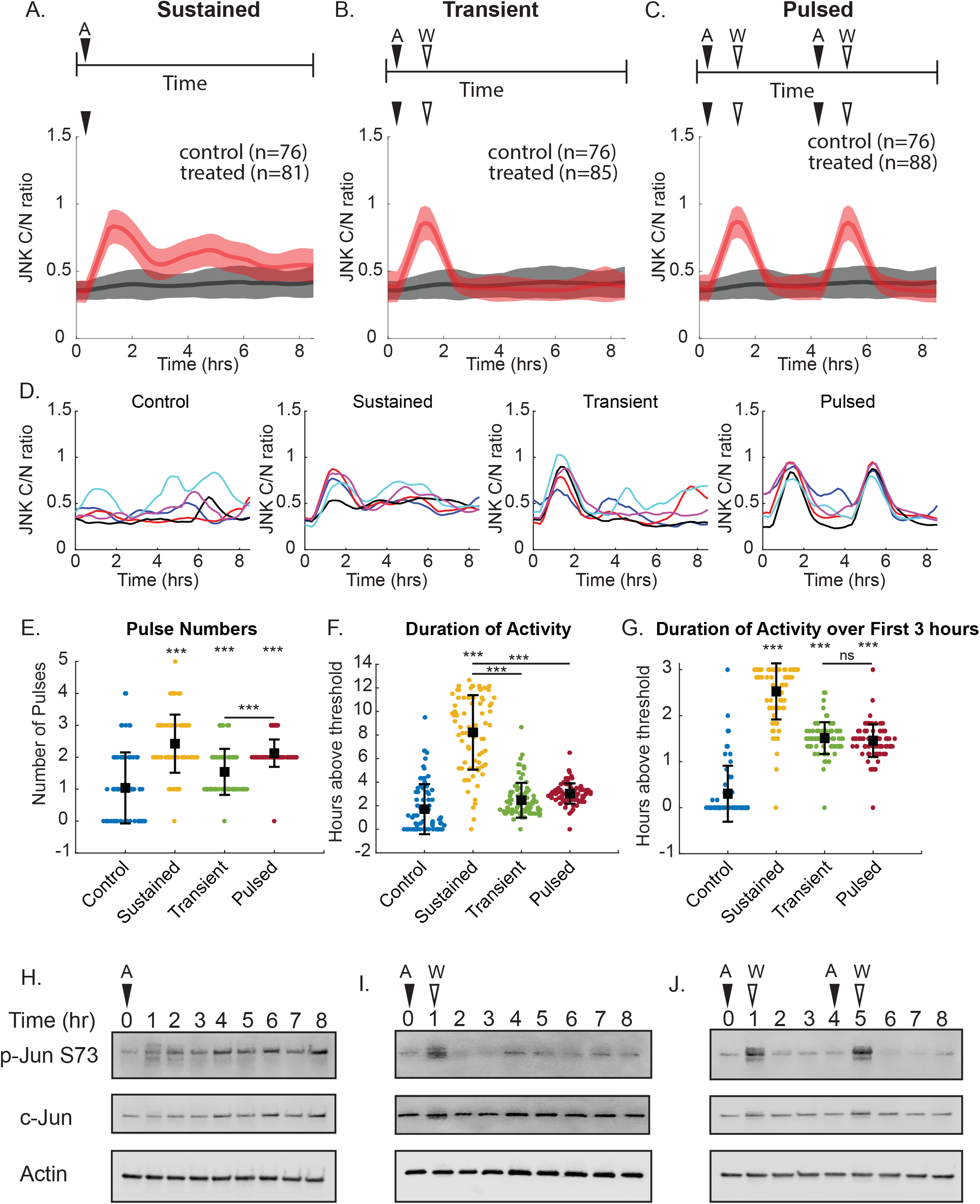
Anisomycin dosing scheme can enrich in specific dynamic patterns. **A-C)** Average JNK C/N ratios of cells treated with sustained **(A)**, transient **(B)**, or pulsed **(C)** anisomycin. Treated cells shown in red versus DMSO vehicle control in black. Thick lines represent the mean response with shaded areas equal to the standard deviation. n is the number of cells analyzed. **D)** Representative traces of five individual cells from each treatment and control condition. **E)** Pulse numbers for each individual cell over the duration of the imaging. Bars shown mean±SD. ***p<0.001 as assessed by ANOVA. **F)** Total duration of JNK activity over the 8.5 hour experiment. Bars show mean±SD. ***p<0.001 as assessed by ANOVA. **G)** Duration of JNK activity over the first 3 hours. Bars show the mean±SD. ***p<0.001 as assessed by ANOVA. Ns=not significant. **H-I)** Western blot images of phosphorylated c-Jun (p-Jun S73), total c-Jun, and β-actin in response to sustained **(H)**, transient **(I)**, or pulsed **(J)** anisomycin treatment confirming phosphorylation patterns of c-Jun mirror KTR traces.

To confirm that JNKKTR dynamics were reflective of endogenous transcription factor activation we performed western blot analysis for phosphorylation of c-Jun at serine 73 (Figure 2H-J). Consistent with the KTR measurements, sustained activation of JNK was characterized by prolonged phosphorylation of c-Jun over the observed time points, beginning as early as one-hour post-stimulation and maintaining high levels up to eight hours (Figure 2H). In the case of transient JNK activation, the Western blot analysis revealed a rapid increase in c-Jun phosphorylation at serine 73, peaking at approximately one-hour post-stimulation, followed by a sharp decline as the pathway returned to baseline activity (Figure 2I). Pulsed JNK activation exhibited periodic phosphorylation of c-Jun at serine 73 (Figure 2J). Studies of the tumor suppressor p53 have shown that mRNA stability mediates gene expression patterns in response to pulsatile p53 and these predicted dynamics can be modeled using ordinary differential equations (ODE). We therefore wanted to determine whether we could similarly predict JNK dependent gene expression patterns using these modeling approaches.

### ODE Model of JNK Signaling Predicts Dynamic Gene Expression Patterns

To predict how JNK dynamics may influence gene expression patterns we employed a simple ODE model based on known regulatory factors within the JNK signaling cascade (Figure 3A). This includes negative feedback from the dual specificity phosphatase DUSP1, which is a transcriptional target of c-Jun. Estimates of protein and mRNA decay rates for the model were based upon literature evidence^25-27^. The model incorporated a linear pathway of JNK activation and a step function to simulate the stimulus. Using this model, we could effectively simulate a sustained (Figure 3B), transient (Figure 3C), or pulsed stimulus (Figure 3D) and predict active JNK and phosphorylated c-Jun levels. The predicted output of active JNK aligned well with our experimental measures of single cell JNK activation (Supplemental Figure 2).

**Figure 3.**
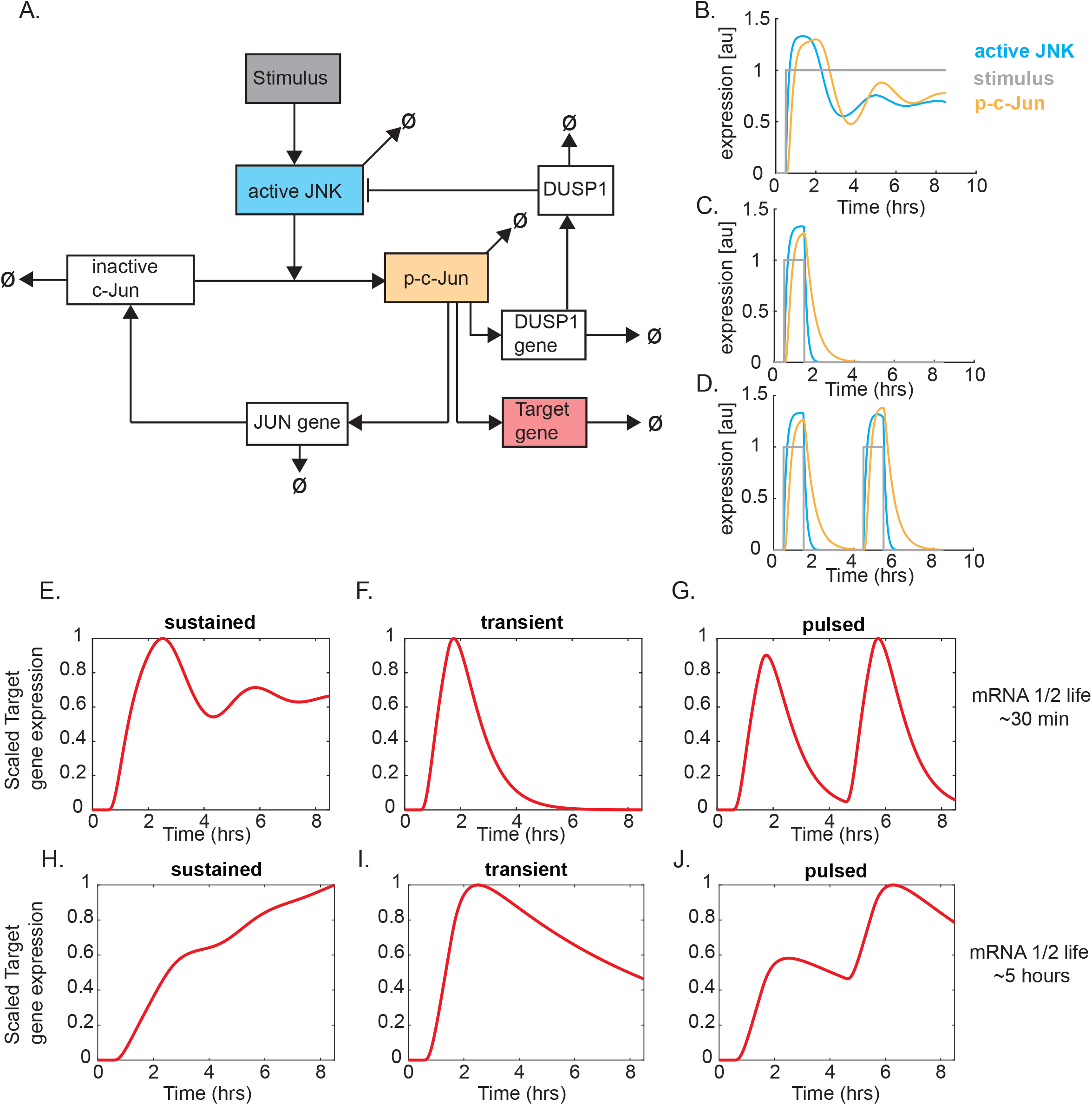
ODE model of JNK signaling predicts dynamic gene expression patterns. **A)** Schematic representation of ODE model of JNK signaling with arrows representing production terms and Ø representing degradation. **B-D)** Simulations of active JNK (blue), phosphorylated c-Jun (orange), and stimulus (gray) for sustained **(B)**, transient (**C)**, or pulsed **(D)** stimuli. **E-J)** Simulated gene expression patterns of target genes for sustained, transient, or pulsed JNK activation with either a short or long-lived mRNA. Data is scaled from 0-1.

The regulation of target gene expression was built on established principles from other dynamic systems, such as p53 and NF-κB^6,12^, where mRNA stability plays a crucial role in determining the timing and magnitude of gene expression. By coupling our model with a downstream transcriptional output, we were able to predict gene expression profiles for short-lived (half-lives=30 minutes) and long-lived (half-lives=5 hours) mRNAs under sustained, transient, and pulsed JNK activation (Figure 3E-J). Specifically, we hypothesized that short-lived mRNAs would exhibit rapid changes in expression, closely mirroring the JNK activation pattern (Figure 3E-G), while long-lived mRNAs would show more integrated responses, reflecting cumulative JNK activity over time (Figure 3H-J). Based on these predicted gene expression patterns, we wanted to determine what patterns would emerge biologically and whether the model could accurately predict these dynamic patterns.

### RNA Sequencing Reveals Gene Expression Dynamics Driven by JNK Activation Patterns

In order to experimentally determine whether the predicted gene expression patterns emerged in cells exhibiting specific JNK dynamics, we treated cells with our dosing regimen as before and collected mRNA at 0, 2-, 4-, 6-, or 8-hours post-treatment. These time points specifically lagged peak JNK activation from our KTR data (Figure 4A) and c-Jun phosphorylation (Figure 2H-J) by 1 hour as previous studies in p53 demonstrated peak gene expression lagging transcription factor activation by approximately 1 hour. As the first pulse of JNK is largely identical between pulsed and transient conditions (Figure 2G), we used the same RNA for these conditions at time points 2 and 4. In addition to the dynamic treatment groups, we also included an additional group which received sustained anisomycin treatment in the presence of the JNK inhibitor tanzisertib (10 µM). Preliminary analysis of the data identified five samples that exhibited poor read quality and were discarded from final analysis. Principal component analysis (PCA) was conducted to visualize the variance in gene expression profiles under sustained, transient, and pulsed JNK activation conditions (Figure 4B). The PCA plot revealed distinct clustering of samples corresponding to each activation pattern, with PC1 and PC2 seeming to correlate with time and JNK activation respectively.

**Figure 4.**
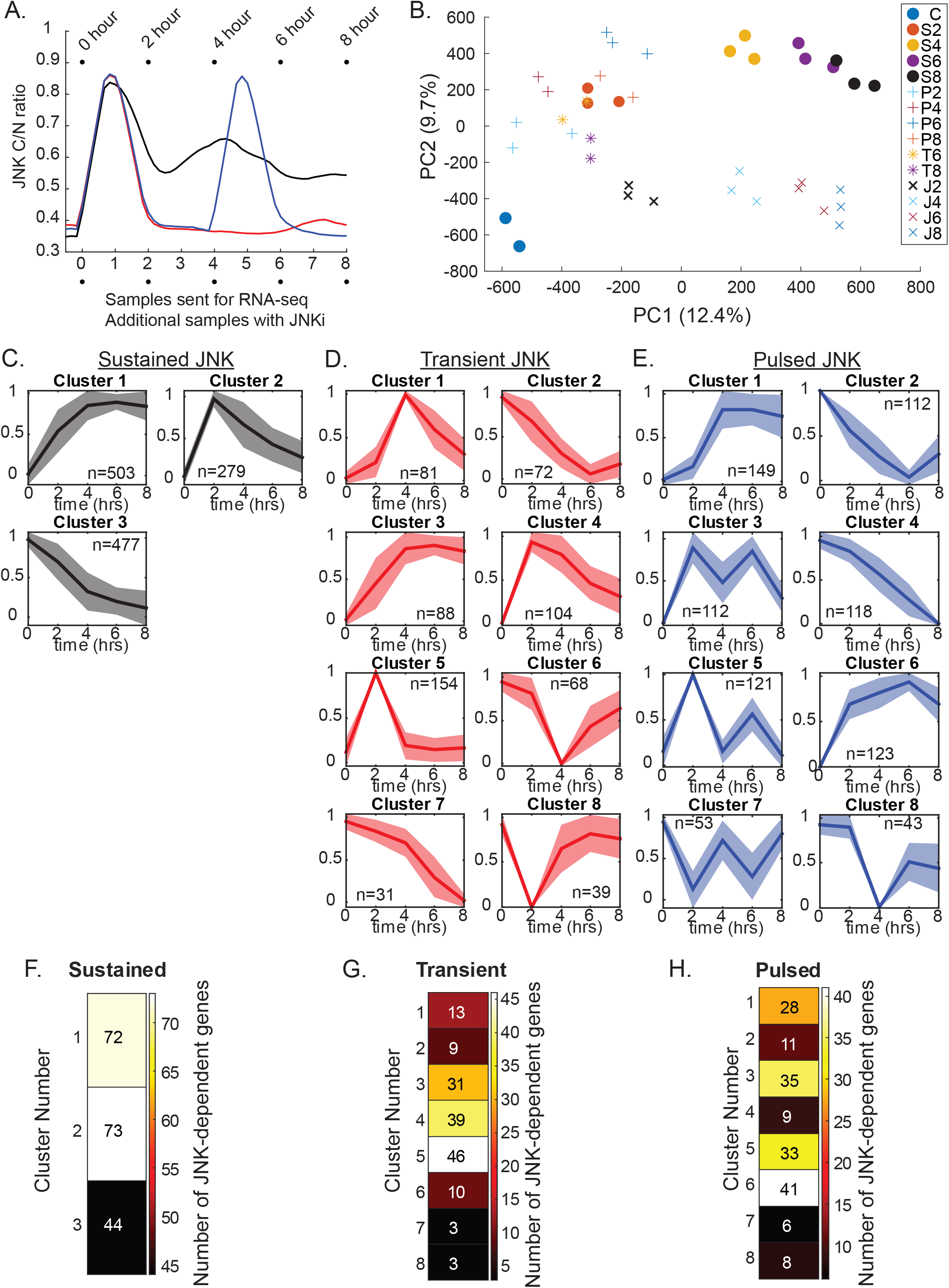
RNA-seq identifies gene expression clusters unique for specific patterns of JNK activation. **A)** Schematic highlighting the timepoints where mRNA was collected. Data traces are mean traces from Figure 2A-C used to highlight relationship of mRNA timepoints to JNK activity. **B)** Principal Component Analysis (PCA) of RNA-seq timepoints and samples. C, ctl; S, sustained; T, transient; P, pulse. Numbers represent time point. **C-E)** K-means clusters of gene expression patterns for sustained **(C)**, transient **(D)**, or pulsed **(E)** JNK activity. Data shown as the mean±SD of scaled expression data. N represents the number of genes within each cluster. **F-H)** Number of JNK-dependent genes identified within sustained **(F)**, transient **(G)**, or pulsed **(H)** clusters.

To identify dynamic gene expression patterns across JNK dynamics, we first identified differentially expressed genes (DEGs) across multiple time points (2, 4, 6, and 8 hours post-treatment) for each activation pattern. If a gene was significantly up- or down-regulated at any of the time points it was included for subsequent analysis. Not surprisingly, the majority of differentially expressed genes were shared across conditions (Supplemental Figure 3A). Fold change differences were scaled using Min-Max normalization to avoid clustering genes based on absolute differences in expression, followed by k-means clustering to group genes exhibiting similar expression dynamics over time (Supplemental Table 1). The clustering analysis revealed distinct gene expression profiles corresponding to each JNK activation pattern (Figure 4C-4E). Under sustained JNK activation, we observed predominantly sustained gene expression clusters, with a few transient clusters that peaked and then declined. Transient JNK activation induced clusters that showed rapid peaks in expression (Figure 4D, clusters 4 and 5), followed by a swift return to baseline. However, other clusters exhibited sustained gene expression (Figure 4D, cluster 3). Pulsed activation of JNK resulted in a few clusters that showed pulsatile gene expression to varying degrees (Figure 4E, clusters 3 and 5). However, as before, other clusters showed more complex dynamics. Interestingly, many of the genes within a given cluster also correlated strongly with other clusters when the dynamics of JNK changed (Supplemental Figure 3B).

As anisomycin can activate additional pathways beyond JNK, we wanted to validate which clusters contained seemingly JNK-dependent genes. To identify these genes, we compared differentially expressed genes (log2 fold change greater than 1) between our sustained condition and our JNKi treatment group which received both anisomycin and a JNK inhibitor. These JNK-dependent genes were then identified based on which cluster they occupied across conditions (Figure 4F-H). As anticipated based on c-Jun’s role as a transcriptional activator most JNK-dependent genes appeared within the upregulated gene clusters. In the case of transient JNK activation, these genes preferentially fell within clusters 3, 4, and 5 (Figure 4G) whereas in response to pulsed activation JNK-dependent genes appeared in clusters 1, 3, 5, and 6 (Figure 4H). Previous studies of p53 dynamics have proposed that diversification of gene expression patterns allows for potential differential regulation of specific pathways over time^12^. We hypothesized that perhaps JNK dynamics facilitated a similar process.

### Dynamic gene clusters are enriched in specific pathways

To explore the potential biological functions of these gene expression clusters, we performed functional enrichment analysis using the ShinyGO v0.82^28^. Only a subset of gene clusters showed enrichment of specific KEGG pathways; however, we did observe that many gene clusters showed enrichment in distinct pathways. For example, sustained clusters 1 exhibited enrichment in a number of apoptosis or cell death associated proteins (Figure 5A), including several pro-apoptotic factors such as *PMAIP1* (Noxa) and *BAK1* (Bak) suggesting prolonged activation of JNK may favor accumulation of these pro-apoptotic signals. In contrast, sustained cluster 2, which exhibits only a transient increase in gene expression, was enriched in several inflammatory signaling pathways, NF-kB response, and MAPK-associated responses (Figure 5B). Interestingly, as many of the genes identified across dynamics are shared similar enriched pathways were found across sustained, pulsed, or transient clusters. However, these exhibited different expression dynamics depending on the pattern of JNK activation.

**Figure 5.**
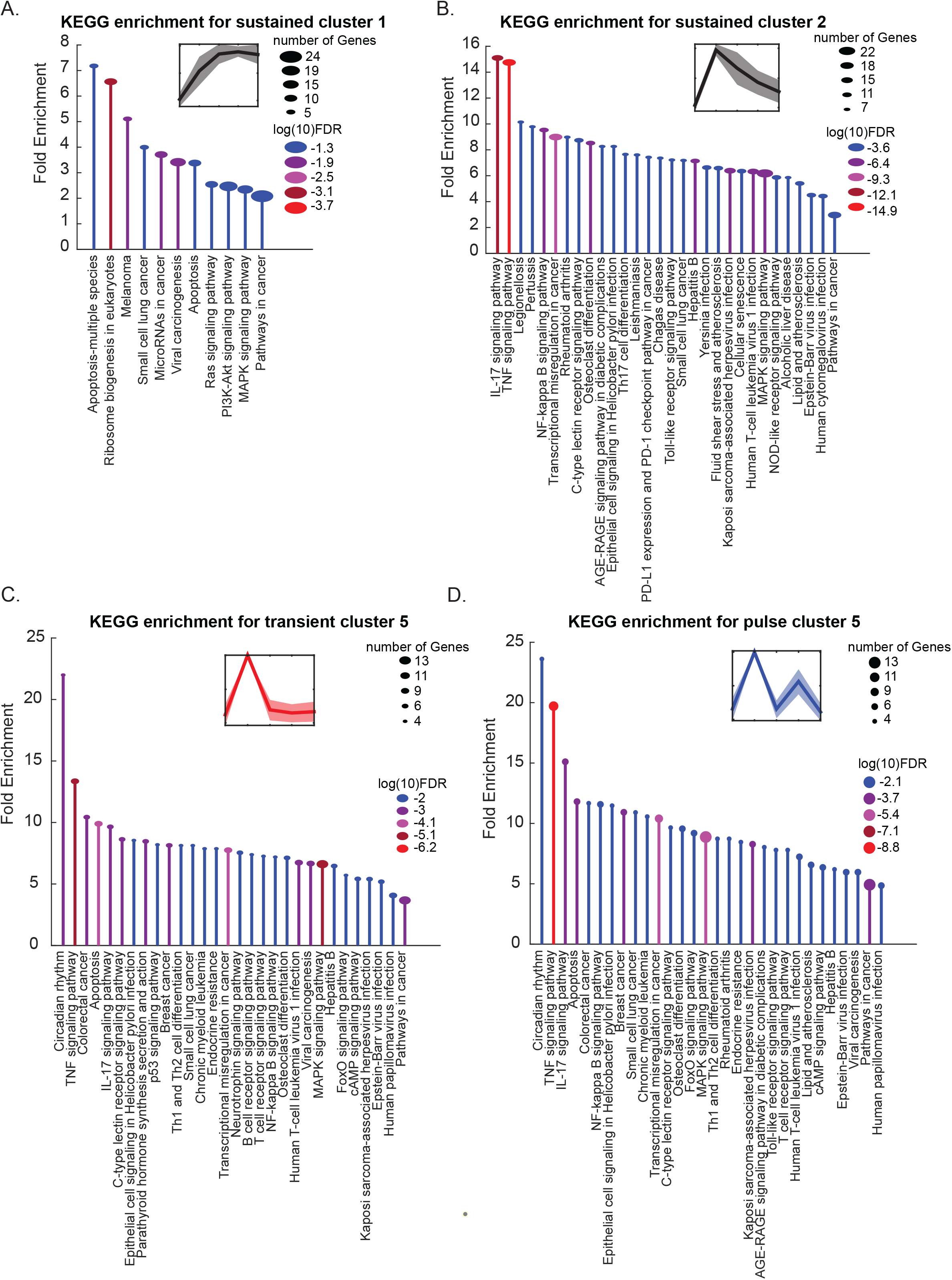
Gene expression clusters are associated with unique cellular signaling pathways. **A-D)** Stem plots for enriched KEGG pathways for different dynamic clusters. **A)** Sustained cluster 1. **B)** Sustained cluster 2. **C)** Transient cluster 5. **D)** Pulsed cluster 5. Color of each stem correlates to the log10 FDR value with the ball size correlating to the number of genes identified. KEGG pathway names are plotted below each stem. Data generated using ShinyGO 0.82 with plots prepared using Matlab.

For example, sustained cluster 1 (Figure 5A), transient cluster 5 (Figure 5C), and pulsed cluster 5 (Figure 5D) all showed enrichment of genes associated with apoptosis but their expression patterns were distinct. Sustained cluster 1 showed sustained expression over time, while transient cluster 5 only showed a transient increase in expression. Pulsed cluster 5 exhibited two pulses of expression. These findings suggested that perhaps variable JNK dynamics would be sufficient to induce different cell fate outcomes in response to the same stimulus. We hypothesized that given the sustained upregulation or repeated pulses of apoptotic genes seen with sustained or pulsed JNK expression we would observe increased cell death under these conditions. However, we observed no increase in sub-G1 cells as assessed by flow cytometry, though we did see a slight increase in G2 accumulation in response to sustained or pulsed JNK activation (Supplemental Figure 4). This suggests additional mechanisms may be required for induction of cell death. Our ODE model predicts that these dynamics should be dictated by the mRNA stability of the specific target genes and we wanted to examine whether this was indeed the major regulatory mechanism involved.

### mRNA stability explains dynamics of select clusters

To examine whether mRNA stability was predictive of our observed dynamic clusters, we compared the experimentally-derived gene expression clusters with the predicted patterns from our ODE model with variable mRNA half-lives (Figure 6A-C). Specifically, we focused strictly on the upregulated gene clusters based upon our model assumptions of c-Jun as a transcriptional activator and consistent with the majority of JNK-dependent genes residing within upregulated clusters (Figure 4F-H). The results suggest that certain clusters may be regulated by mRNA stability; however, this does not appear to be the only potential mechanism. For example, neither sustained cluster 1 nor cluster 2 were explained by mRNA stability alone, though cluster 1 did appear to align more closely to long-lived mRNA while cluster 2 was more similar to short-lived mRNA (Figure 6A). In contrast, transient clusters 4 and 5 appear to be largely explained by differences in mRNA stability while transient clusters 1 and 3 deviate strongly from the ODE model (Figure 6B). For the pulsed clusters, the majority did not align well with the ODE model, though clusters 3 and 5 do appear to correlate with predictions of short-lived mRNA (Figure 6C).

**Figure 6.**
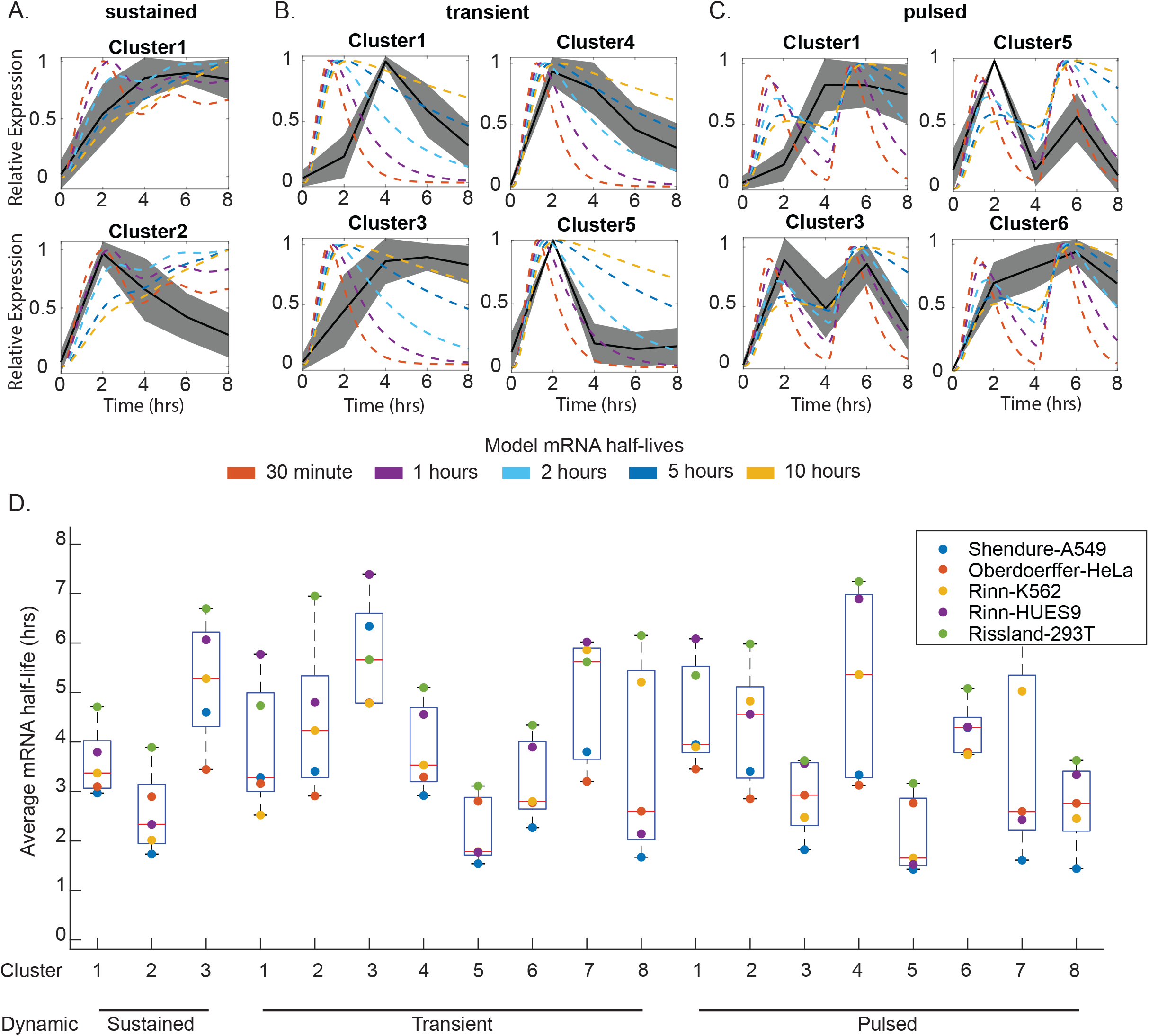
mRNA stability contributes to gene expression clusters driven by JNK activation. **A-C)** Predicted scaled mRNA expression from our ODE model of JNK signaling were aligned to experimentally measured gene expression clusters for sustained **(A)**, transient **(B)**, or pulsed **(C)** JNK activity. Colored lines represent predicted mRNA expression for variable mRNA half-lives with black lines showing experimentally measured gene expression dynamics. **D)** Box plots showing average mRNA half-lives from multiple studies for genes within each identified cluster.

To determine whether our predictions aligned with measured mRNA half-lives, we examined multiple published datasets of global mRNA stability^29-32^. From these studies we collected mRNA half-life measurements and cross-referenced these genes with the genes within each of our experimental clusters. For each study, we calculated the average mRNA half-life of genes within each cluster. One limitation of this approach is that not all genes appear in every dataset. However, based on this analysis, we found that many of the predictions aligned with experimental measurements. Genes within sustained cluster 2 do have shorter mRNA half-lives than genes from sustained cluster 1 in these studies. Similarly transient clusters 4 and 5 corresponded to longer lived and short-lived mRNA respectively. Pulsed clusters 3 and 5 also corresponded to short-lived mRNA as predicted. While these clusters were supportive of mRNA stability as a driver further investigation is needed to identify other major determinants of the observed dynamics for other clusters.

## Discussion

The JNK signaling pathway plays a pivotal role in cellular responses by balancing the induction of cell survival and cell death^16,33^. The temporal dynamics of JNK activation have previously been shown to promote distinct cellular fates across multiple stimuli including UV light, oxidative stress, and endoplasmic reticulum stress^16,17,19-21^. Within our studies, we have found that these dynamics vary not only across various stimuli, but also within individual cells exposed to the same stimulus, as cells exhibit variability in the duration or pulses of JNK activation over time. This variability within cells likely stems from variable levels of positive and negative regulators, similar to the findings from the Aoki lab, where variations in p38 and DUSP1 levels contributed to heterogeneity in JNK activation^19^. However, how these dynamics from previous studies ultimately contribute to downstream gene expression patterns has been unclear.

Our study demonstrates that variations in the duration or pulse numbers of JNK activity can elicit unique gene expression profiles. Similar to previous work examining p53 and NF-kB, these dynamics appear to be influenced in part by mRNA stability^6,12,34^. However, mRNA stability alone appears insufficient to explain all the observed gene expression patterns. One possibility is that some of the observed gene expression patterns may rely on transcriptional cross-talk by additional transcription factors that may or may not be regulated by JNK. Consistent with this possibility, many of the observed genes were associated with NF-kB activity. Previous studies looking at the decoding of signals through pathogen-associated molecular patterns (PAMPs) have suggested an array of combinatorial networks that drive gene expression patterns^3^. Included among these identified networks is a coherent feedforward loop regulated by both NF-kB and AP-1, as well as coordinated cross-talk between NF-kB and p38^3^. It is well established that anisomycin activates both JNK and p38, potentially providing another route for coordinated regulation of some of our observed gene expression patterns. In order to examine whether these other clusters rely on coordination between these pathways, a more in-depth characterization of cis-regulatory elements in the promoters of these genes, as well as characterization of NF-kB and p38 dynamics are needed, which we aim to undertake in future studies. One additional mechanism that may explain clusters that do not behave as expected based solely on mRNA stability is the potential inhibitory phosphorylation of c-Jun. Recent work suggests that prolonged JNK activation hinders AP-1 activity by phosphorylation of c-Jun at Threonine 91 and 93^35^. This mechanism may explain why sustained activation of JNK does not drive a continuous increase in gene expression as we observed a gene cluster that exhibited transient upregulation with sustained JNK activation.

Similar to p53 and NF-kB expression patterns^6,7,12^, JNK dynamics appear to drive differential expression patterns of genes involved in specific pathways, such as cell death and inflammation-associated genes. These patterns may allow cells to differentially interpret stimuli. For example, genes that are rapidly and transiently induced may allow cells to reset and prepare for an additional stimulus, whereas genes that appear to accumulate over time, including cell death genes, may allow cells to integrate sustained signals until a critical threshold is met to induce cell death. While we did not observe any major distinctions in cell death by flow cytometry, we did observe slight variations in cell cycle arrest depending on the dynamics of JNK activation. Future studies examining how the frequency of JNK activation contributes to these patterns may provide further elucidation of differential functions of these dynamics.

Mechanistically, these studies point to a highly conserved mechanism of regulating gene expression patterns through a combination of upstream signaling dynamics and mRNA stability. However, unlike p53 and NF-kB, JNK activation dynamics may have additional effects beyond solely transcriptional regulation,includingregulation of biological function through phosphorylation of additional substrates and through localization to specific subcellular compartments^36-39^. Further study is needed to understand how these additional functions coordinate within our basic model of transcriptional regulation. Clinically, the targeting of protein dynamics has proven difficult, particularly for transcription factors; however, kinase targeting has been far more successful. As a result, understanding the function of these dynamics and potential avenues to manipulate them may have significant impacts on potential treatment of human disease.

## Materials and Methods

### Cell lines and culture

hTERT-RPE1 cells were originally purchased through ATCC (CRL-4000) and were grown in DMEM/F-12 media (Gibco:Hygromycin B Solution (50 mg/mL), Catalog Number 10687010.) supplemented with 10% fetal bovine serum (FBS; Corning.), 100 U/mL penicillin, and 100 µg/mL streptomycin. The cultures were maintained in a humidified incubator at 37°C with 5% CO_2_. RPE1-hTERT cells expressing H2B-mCerulean and JNKKTR-mClover were generated via lentiviral transduction and selection with hygromycin and puromycin respectively. Following selection, single cell clones were generated by serial dilution and screened before imaging. All imaging studies were performed using clonally derived cells to minimize impacts due to genetic differences.

### Induction of JNK dynamics

To induce JNK activation, cells were treated with 50 ng/ml of anisomycin. For sustained activation of JNK, cells received a single treatment with anisomycin. To induce transient activation cells were treated with anisomycin for 1 hour followed by three washes with DMEM/F12 growth media and then placed into fresh media. Pulsed activation of JNK was induced by introducing a second anisomycin treatment at 4 hours followed by an additional washout 1 hour later as performed for transient conditions. Additional treatments were performed using sorbitol (Fisher, BP439-500), tunicamycin (Fisher, 35-161-0) thapsigargin (Fisher, 11-381), H_2_O_2_(Sigma-Aldrich, H1009-100ML), TGF-β1 (Fisher, GF346), and TNF (Fisher, 501615266) at the indicated concentrations.

### Live-cell microscopy

Live-cell imaging was performed using Nikon Ti2 widefield fluorescence microscope equipped with Tokai-Hit environmental chamber for gas and humidity control. Images were captured every 10 minutes using a Plan Apo λD 20X/0.8 NA objective. Excitation of fluorescent proteins (H2B-Cerulean and JNKKTR-mClover) was performed using a Spectra III light engine (CFP, 440 nm; YFP, 514 nm) with LED intensity set at 25%. CFP light was passed through a CFP / YFP / mCherry - Pinkel Triple excitation filter and 474/29 excitation filter. YFP excitation light passed through the same Pinkel Triple filter followed by 544/24 emission filter. Images were captured using an OrcaFusion BT camera on 16-bit image setting to capture H2B-mCerulean (200 ms exposure) and JNKKTR-mClover (300 ms exposure). Images were exported as individual tiff images for each time point and position for subsequent analysis.

### Quantification of Kinase C/N Ratio

Quantification of live-cell microscopy was performed using MATLAB p53Cinema[64]. Briefly, cells were segmented and tracked using captured H2B-mCerulean images with a minimum of 50 individual cells tracked per condition. Identified centroids were used to measure mean fluorescence intensity of JNKKTR mClover within the indicated regions of interest. Cytoplasmic-to-nuclear ratios (C/N ratio) were quantified by expanding the identified nucleus by 3 pixels to generate a cytoplasmic ring to approximate cytoplasmic fluorescence as has been used in other studies^18^. The ratio between the cytoplasmic ring and nuclear measurements gave the C/N ratio. For quantification of pulse numbers, we used the findpeaks function in Matlab with the assignment of a minimal peak threshold of 50% above average JNK activity at baseline to avoid spurious peak detection due to noise in individual cells. Quantification of overall durations of JNK activity was performed by calculating the duration of JNK activity above this same threshold.

### Western Blot Analysis

Cells were grown on 6 cm plates and treated according to induction (sustained, transient, or pulsed) method. After treatment cells were washed with 1× PBS after aspirating the culture medium. Cells were collected by scraping in 1 ml PBS and transferred to microcentrifuge tubes. An additional 1ml of PBS was used to collect remaining cells. The collected cells were centrifuged at 13,000 rpm for 5 minutes, lysed in lysis buffer (ThermoFisher, 89900) containing protease and phosphatase inhibitors (ThermoFisher, 78440), and incubated on ice for 30 minutes with agitation. Insoluble debris was removed by centrifugation at 13,000 rpm for 10 minutes at 4°C. Protein concentration was determined by Bradford assay (Biorad, 5000006) using a standard curve of known BSA concentration.

Proteins (20 µg per lane) were separated on 4-20% Tris-Glycine gels (Biorad, 4561096) and transferred to PVDF membranes (Millipore, IPFL00010). Membranes were blocked in Licor Blocking Buffer (Licor, 927-70001) for 1 hour and incubated overnight at 4°C with primary antibodies specific to the target proteins. Primary antibodies include c-Jun (CST#9165; 1:1000), Phospho-c-Jun serine 73 (CST#3270, 1:1000) and B-actin (CST#4970,1:3000). After washing three times with 0.1% PBST, membranes were incubated with IRDye secondary antibodies (Licor) at 1:5,000 dilution for 1 hour, washed again, and imaged using a Licor Odyssey Fc imaging system. All experiments were performed using three biological replicates.

### RNA sequencing

Cells were grown on 6 cm plates and treated according to induction method (sustained, transient, or pulsed JNK activation). Additionally, for RNA-sequencing we included an additional group which received treatment with the JNK inhibitor tanzisertib (10 uM) in addition to sustained anisomycin treatment. Total mRNA was isolated using the Qiagen RNeasy Mini Kit (Qiagen, 74106) at the indicated time points (0, 2, 4, 6, or 8-hours post-treatment). RNA was provided to the SDSU Genomics Sequencing Core Facility for quality assessment, library preparation, and sequencing. Libraries were prepared using the Illumina TrueSeq mRNA and 45 samples prepared for NExtSeq500 (1×75bp) High output reads. Sequencing was performed using 3 runs to get sufficient reads for all samples and performed by the SDSU Genomics Sequencing Core Facility. All bioinformatic analyses were done using CLC Genomics Workbench (vs 24.0). Trimmed reads were mapped to the human genome vs hg38 as reference using a minimum length fraction of 0.8 and a minimum similarity fraction of 0.8. Differential expression was calculated on a gene basis using an absolute fold change cut-off of 2 and a FDR adjusted p-value of 0.05.

### Ordinary Differential Equation (ODE) modeling of JNK signaling

In order to model predicted gene expression dynamics, we designed an ODE model using Matlab based on 7 species. These 7 species and the equations used are highlighted below. . To model JNK activation we used a constant (c) which we varied between 0 and 1 as a step stimulus. Estimates of rate parameters were pulled from literature or based upon p53 dynamic models^10^. For modeling variations in mRNA stability, we varied the mRNA decay rate (a_m).

| Equation Species | Species | Equation |
| --- | --- | --- |
| Y(1) | Inactive c-Jun protein | $dy(1) = b_j[y(3)] - a_j[y(1)] - b_{jp}[y(1)] \frac{y(5)^n}{y(5)^n + T^n}$ |
| Y(2) | Phosphorylated c-Jun protein | $dy(2) = b_{jp}[y(1)] \frac{y(5)^n}{y(5)^n + T^n} - a_{junp}[y(2)]$ |
| Y(3) | c-Jun mRNA | $dy(3) = b_{jmi} + b_{mj}[y(2)] - a_{jm}[y(3)]$ |
| **Y(4)** | Target gene mRNA | $dy(4) = b_t + b_{tj}[y(2)] - a_m[y(4)]$ |
| **Y(5)** | Active JNK | $dy(5) = b_{jnkp}[c] - a_{jnkp}[y(5)] - ad_{jnkp}[y(5)] \frac{y(7)^n}{y(7)^n + Td^n}$ |
| **Y(6)** | DUSP1 mRNA | $dy(6) = b_{md}[y(2)] - a_{dusp}[y(6)]$ |
| **Y(7)** | DUSP1 protein | $dy(7) = b_{dusp}[y(6)] - a_{dp}[y(7)]$ |

| Parameter | Model Value |
| --- | --- |
| b_jmi | 1 |
| b_mj | 1.5 |
| a_jm | 2.1 |
| n | 4 |
| b_jp | 3 |
| T | 1 |
| a_junp | 2.1 |
| b_j | 3 |
| a_j | 0.5 |
| b_t | 1 |
| b_md | 0.25 |
| b_tj | 3 |
| a_m | 0.14 |
| b_jnkp | 10 |
| a_jnkp | 7.5 |
| c | 0 or 1 |
| a_dusp | 2.1 |
| ad_jnkp | 50 |
| b_dusp | 1 |
| a_dp | 0.693 |
| Td | 0.2 |

### K-means clustering

K-means clustering of differentially expressed genes was performed using Matlab. Briefly, differential expression analysis from CLC was imported into Matlab and significantly differentially expressed genes identified based on an FDR-adjusted p-Value <0.05, an absolute Log_2_Fold Change >1, Max Group Mean >1, and restricted to protein coding transcripts. Data was organized based on treatment condition and timepoint then rescaled between 0 and 1. K-means clustering was then performed on each condition (sustained, pulsed, or transient) using the k-means function with the default seed. Selection of the cluster numbers was determined using combinations of silhouette score, elbow method, and minimizing variations in cluster size.

### Cell Cycle Analysis

3.5 × 10^5^ hTERT-RPE1 cells were plated onto 6-cm plates (Genesee) and allowed to grow for 24 h. At the time of treatment, media were replaced with fresh media and treated cells received anisomycin treatments as previously outlined. After 24 or 48 h, media were collected and placed into labeled tubes. Cells were trypsinized and added to the collection tube containing the original media. Cells were spun down at 400 *g* for 4 min and washed twice with PBS to remove residual media. Cell pellets were re-suspended in 300 μl PBS and passed through a 45 μm cell strainer to remove clumps. Cells were then fixed by the addition of 700 μl cold ethanol dropwise. Fixed cells were stored at −20°C until staining. Prior to staining, cells were spun down at 750 *g* for 5 min and washed twice with PBS to remove ethanol. 1 × 10^6^ cells were then re-suspended in 250 μl PBS + 10 μl propidium iodide and 5 μl RNaseA. Cells were incubated at 37°C for 30 min, then brought to 500 μl final volume by PBS and passed through a 45 μm cell strainer prior to analysis on a BD Accuri C6. Single cells were identified based on initial gating on forward and side scatter measurements followed by PI-A and PI-H gating. Gates for sub-G1, G1, S, and G2/M phases were manually annotated to histograms of the PI stain. Data for cell cycle distributions were presented as the mean ± SEM of three biological replicates.

## Data Access and Availability

Single cell KTR traces used within this study have been uploaded to Mendeley Data with unique identifiers. The basic JNK signaling model has also been uploaded to Mendeley Data. RNA-seq data was uploaded to GEO.

JNK KTR in response to multiple stimuli (10.17632/4s278fpzp4.1)

Manipulation of JNK dynamics by Anisomycin (10.17632/sjkzrs3sfz.1)

JNK model of dynamic gene regulation (10.17632/h828r74cjt.1)

RNA-seq data: GEO (accession provided once dataset approved)

## Acknowledgements

This material is based upon work conducted using the South Dakota State University Functional Genomics Core Facility (RRID:SCR_023786) supported in part by the National Science Foundation/EPSCoR Grant No. 0091948, the South Dakota Agricultural Experiment Station, and by the State of South Dakota. This material is based upon work conducted using the South Dakota State University Genomics Sequencing Core Facility (RRID:SCR_023959) which is supported in part by the South Dakota Agricultural Experiment Station. Research reported in this publication was supported by the National Institute of General Medical Sciences of the National Institutes of Health under Award Number P20GM135008. The content is solely the responsibility of the authors and does not necessarily represent the official views of the National Institutes of Health. Funding for this research was provided by the South Dakota Board of Reagents Competitive Research Grant program (SA2400045).

## Author contributes

AJ and RLH planned experiments. AJ, RB, AC, and RLH performed and analyzed experimental data. JGH assisted with RNA-sequencing and analysis. AJ, RB, JGH, and RLH wrote and proofread the manuscript.

## Disclosure and competing interests statement

The authors declare no competing interests or disclosures

